# Multilabel multiclass classification of OCT images augmented with age, gender and visual acuity data

**DOI:** 10.1101/316349

**Authors:** Parmita Mehta, Aaron Lee, Cecilia Lee, Magdalena Balazinska, Ariel Rokem

## Abstract

Optical Coherence Tomography (OCT) imaging of the retina is in widespread clinical use to diagnose a wide range of retinal pathologies and several previous studies have used deep learning to create systems that can accurately classify retinal OCT as indicative of one of these pathologies. However, patients often exhibit multiple pathologies concurrently. Here, we designed a novel neural network algorithm that performs multiclass and multilabel classification of retinal images from OCT images in four common retinal pathologies: epiretinal membrane, diabetic macular edema, dry age-related macular degeneration and neovascular age-related macular degeneration. Furthermore, clinicians often also use additional information about the patient for diagnosis. Second contribution of this study is improvement of multiclass, multilabel classification augmented with information about the patient: age, visual acuity and gender. We compared two training strategies: a network pre-trained with ImageNet was used for transfer learning, or the network was trained from randomly initialized weights. Transfer learning does not perform better in this case, because many of the low-level filters are tuned to colors, and the OCT images are monochromatic. Finally, we provide a transparent and interpretable diagnosis by highlighting the regions recognized by the neural network.

## 1. Introduction

Convolutional Neural Networks (CNNs) and other deep neural networks have enabled unprecedented breakthroughs in developing artificial intelligence systems to perform computer-assisted diagnosis based on clinical data and several recent studies demonstrate the ability of these algorithms to leverage large clinical datasets to learb how to classify images as exhibiting a pathology Lee et al. (2016); Choi (2017); Esteva (2017); Kermany et al. (2018). However, most of these studies classify the presence or absence of a single pathology Lee et al. (2016). Even when multiple classes are present in the data, comparisons are usually binary between two different classes of pathologies Esteva (2017) or classify multiple classes of pathology, without accounting for the presence of more than one pathology in the same patient Choi (2017). These classification tasks belie the complexity of clinical data analysis: patients in a clinical population may exhibit several pathologies at the same time or none at all. For example, in the present study, we examined an extraction of 36,150 unique examination with Optical Coherence Tomography (OCT) images from the Electronic Medical Records (EMR) of the Ophthalmology clinic at University of Washington. We found that 24% of patients with any pathology displayed multiple pathologies. OCT uses light to capture *in-vivo* high resolution optical cross sections of retinal tissue. It has become one of the most commonly performed medical imaging procedures, with approximately 30 million OCT scans performed each year world-wide Swanson and Fujimoto (2017). OCT is critical in delineating retinal and choroidal pathologies, and has been shown to be more sensitive in detecting retinal disease than other modalities such as color fundus photography. (PMID: 22347793, 19079147, 26719967). Despite the high-quality information provided by OCT, clinicians often use additional information about the patient like age and visual acuity, along with the OCT scans to aid diagnosis.

Advances in deep learning LeCun et al. (2015) have allowed for significant gains in the ability to classify images and detect objects. This technique typically requires a large dataset, with millions of samples. Transfer learning is among the techniques often used to address the lack of data in a given domain by leveraging data from another domain Bengio and Others (2012). However, a recent study Rokem (2017) demonstrates that we do not need large data corpus to train deep neural networks if the image data is relatively homogeneous, as is the case with OCT images. Most transfer learning is performed on models trained on 1000 class data from ImageNet. ImageNet images are 3 channel (RGB) images and the OCT images are single channel (grayscale). We find this leads to transfer learning not be an effective tool for OCT images. Visualization of the convolution kernels Krizhevsky et al. (2012)show lower level kernels detecting colors in addition to edges and contours. Since the lower layer weights are frozen in transfer learning, these filters result in ‘dead’ features for grayscale OCT images. This can be seen in plateaued learning rates for transfer learning in our experiments.

We train a deep neural network for multilabel multiclass classification. We use patient information like age, gender and visual acuity as features in addition to the pixel data from OCT images. Our model perform 14.37% better on accuracy overall and 113.37% better on exact match compared to a transfer learning model trained on ImageNet data. The addition of age and visual acuity attributes to the model demonstrates the most improvement in accuracy and F1 scores. The addition of gender results in no improvements in three out of four pathologies and decreases the F1 score in case of NVAMD diagnosis.

## 2. Cohort

Our study cohort consisted of an extraction of 2.6 million OCT images linked to clinical datapoints from EMR. Automated extraction of an OCT imaging database was performed and linked to clinical endpoints from the EMR. OCT macula scans were obtained by Heidelberg Spectralis, and each OCT scan was linked to EMR clinical labels extracted from EPIC. Labels from the EMR were then linked to the OCT macular images, and the data was stripped of all protected health identifiers.

### 2.1 Cohort Selection

Patients with Epiretinal membrane (ERM), Diabetic macular edema (DME), Dry age-related macular degeneration (DryAMD) and Neovascular age-related macular degeneration (NVAMD) were chosen; patients with other macular pathology by ICD-9 code were excluded. No images were excluded due to image quality. Age, gender and visual acuity fields were also extracted from the EMR. As most of the macular pathology is concentrated in the foveal region, the decision was made a priori to select the central 11 images from each macular OCT set. Each image was then treated independently and labeled as either normal or containing one of the pathologies enumerated above. The complete data set consists of 36148 unique examinations and 7929 unique patients. Based on the four chosen pathologies, our data set has 11452 unique examinations over 2233 patients. Each examination consists of 11 images, one per slice. Table:1 shows the distribution of the data set over pathologies. 7% of the images exhibit multiple pathologies. DryAMD and NVAMD are mutually exclusive; therefore the patients with multiple pathologies may only exhibit one form of AMD. The images are histogram equalized and center cropped to (299, 299) pixels.

**Table 1:** Pathology distribution over total number of images in dataset.

| Pathology | Number of Images | Number of Unique Examinations |
| --- | --- | --- |
| Epiretinal membrane(ERM) | 24,494 | 2,226 |
| Dry AMD(DryAMD) | 57,709 | 5,246 |
| Neovascular AMD(NVAMD) | 13,165 | 1,196 |
| Diabetic macular edema(DME) | 21,304 | 1,936 |
| Multiple pathologies | 9,306 | 846 |

## 3. Methods

A modified version of the Inception Resnet V2 CNN (Szegedy, 2016) was used as the deep learning model for classification fig. 1^1^. To choose the model architecture we used accuracy on a holdout set and the learning curve. While both Resnet and Inception models show impressive performance on the data set Inception Resnet V2 consistently showed lower training loss and better overall accuracy across classes on the holdout set. In addition, modifications to the models were required for direct training on all the models above. The modifications included changing model parameters to train on grayscale images instead of the RGB images, as well as removing the last softmax layer. The softmax was replaced by sigmoid for assigning labels. We used RMSprop, Tieleman and Hinton (2012) as the optimizer and Multilabel Soft Margin Loss function (sometimes referred to as Sigmoid Cross-entropy loss). This function creates a criterion that optimizes a multilabel one-versus-all loss based on max-entropy, between input *x* and target *y*, where *x* represents k-hot encoding of input image labels.

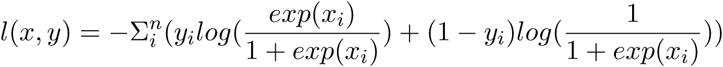

**Figure 1:**
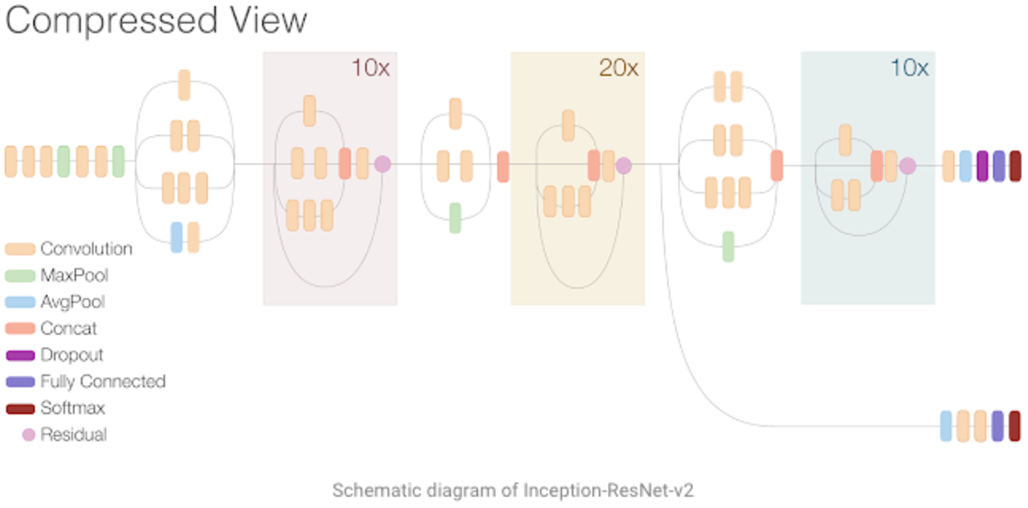
Inception Resnet V2 Architecture

**Figure 2:**
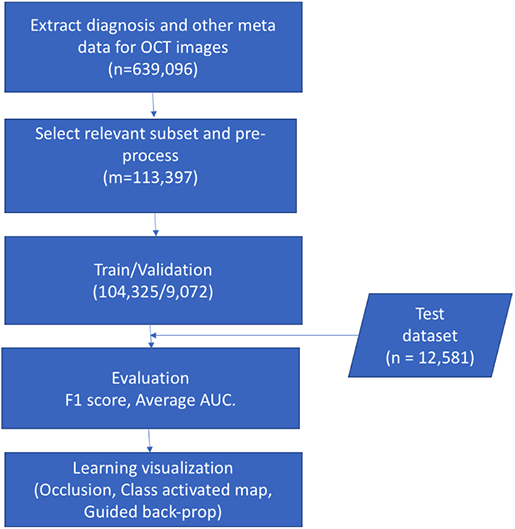
process flow chart

One of the objectives of this study was to compare transfer learning, to learning from a randomly initialized network. Transfer learning experiments were performed on pre-trained Inception Resnet V2 trained on ImageNet data. Pretrained weights were loaded and frozen for the convolutional layers, and the final linear layer was trained to recognize the zero or more of the four disease classes. We used the identical data set and loss function as in training from randomly initialized weights, except that for transfer learning the OCT data with only a single channel was replicated to three channels.

Image data was augmented by applying two transformations. First transformation randomly flipped the image on the horizontal axis, while the second transformation randomly rotated the image between −15°, +15°. The data set was shuffled for every training epoch(iterations through the entire data set). Each epoch consisted of 2,900 iterations. The model was trained until the validation error stopped dropping after one drop of learning rate. Test set was 12,581 images and training/validation set were 104,352/9,072 images respectively. Test and validation sets were chosen to have the same ratio of samples per pathology as the training set. To avoid memorization of individual patient characteristics and overfitting to these characteristics, the test set and training set were seperated at the patient level, as opposed to the exam level. We trained our network utilizing (Pytorch (pyt)) framework on a single Nvidia Titan X GPU, which resulted in batch size being 35. We started with a relatively small learning rate of 1e-5, which was further decreased every 20 epochs by a factor of 0.5. The model was initialized with random weights.

In addition to OCT images, age, gender and visual acuity, (logMAR) figures were extracted from EMR. Age and visual acuity values were retained as floating point numbers while gender was converted to categorical variable (0:male, 1:female). These were then added as additional features directly in the fully connected layer.

## 4. Results

We performed two sets of experiments. The first set of experiments evaluated the utility of transfer learning vs direct learning for multilabel multiclass classification of OCT images. The second set of experiments evaluated the effectiveness of additional information like age, gender and visual acuity on the classification results. All of the models were trained on the same data with identical preprocessing and augmentation techniques.

### 4.1 Evaluation Metrics

Multilabel classification implies finding a model that maps inputs x to binary vectors y (assigning a value of 0 or 1 for each element (label) in y). Evaluation metrics for multilabel classification performance are inherently different from those used in multiclass (or binary) classification, due to the differences in the classification problem. We use the following metrics to measure performance, if T denotes the true set of labels for a given sample and P the predicted set of labels, then:

- Hamming loss: the ratio of the wrong labels to the total number of labels, i.e.

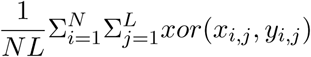

where *y_i,j_* is the target and *x_i,j_* is the prediction.
- Precision, Recall and F1 score: Precision is 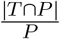, recall is 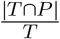 and F1 score is the harmonic mean of precision and recall.
- Exact match: the most strict metric, indicating the percentage of samples that have all their labels classified correctly.

We use average overall accuracy (1 - Hamming loss), exact match and F1 scores for each class as evaluation metrics.

### 4.2 Transfer learning vs Direct learning

Transfer learning has proven to be a highly effective technique, particularly when faced with domains with limited data (Donahue et al., 2014; Yosinski et al., 2014). However, we find transfer learning to not be as effective when using a model trained on ImageNet dataset to build a classifier for OCT data. We achieve much higher accuracy as well as F1 scores, Fig. 3 with the four pathologies on direct training. OCT images are single channel (grayscale) images, while the ImageNet data are color images. The examination of low level features for images with the transfer learning model reveals several dead features. The learning curves for transfer learning vs direct learning show a marked difference. With the same learning rate and learning rate decay schedule for both methods, loss values for transfer learning plateaus. Fig. 4 We tried tuning hyper parameters like learning rate and learning rate decay, but the results for transfer learning did not change. Overall accuracy and exact match for transfer learning was 74.5% and 30.14% compared to 85.23% and 64.3% for direct learning.

**Figure 3:**
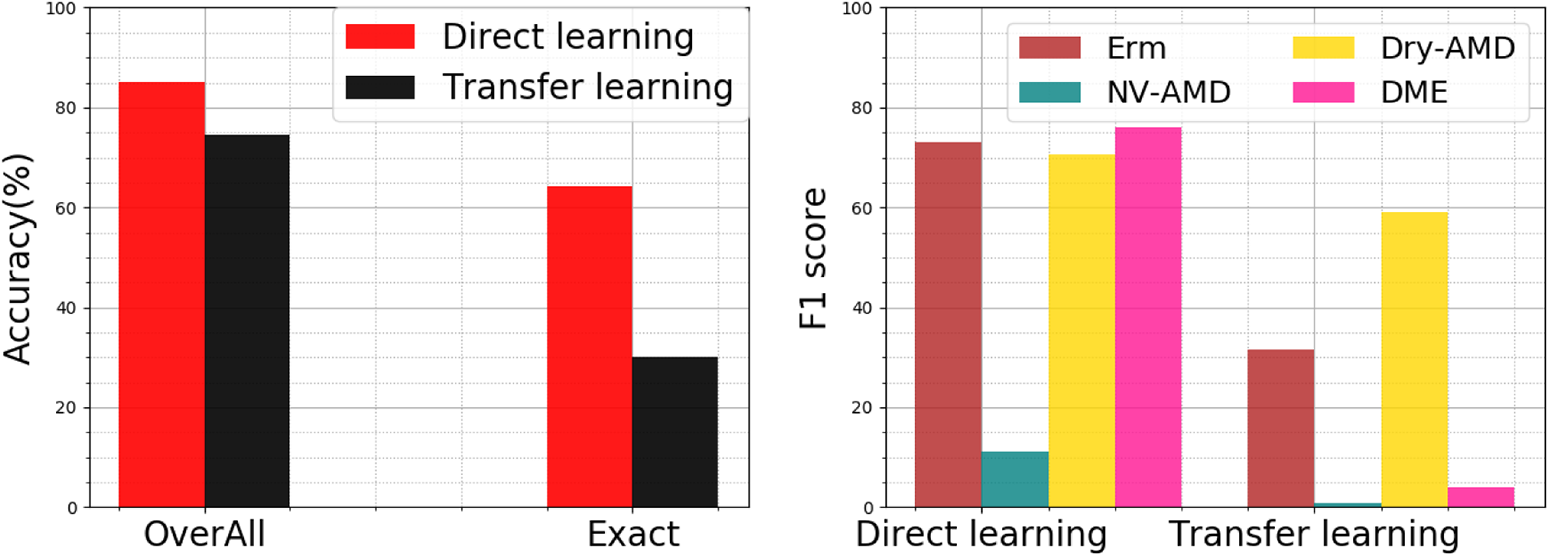
Overall and exact match accuracy (left), F1 score for per pathology (right) using image data for transfer and direct learning.

**Figure 4:**
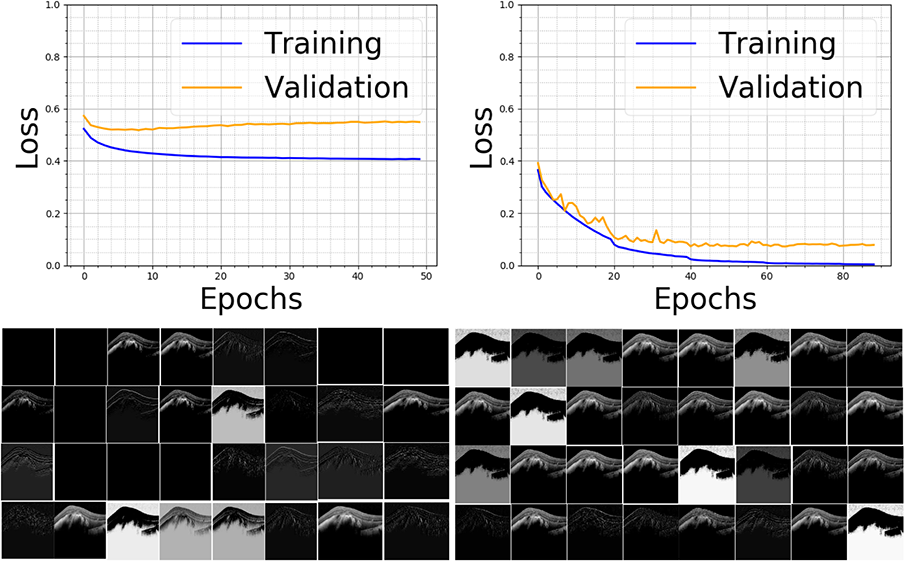
Top row shows learning curves and the bottom row shows the activations from the first layer of the network. The first column is is transfer learning and second column is direct learning. The completely dark patches show dead features.

### 4.3 Classification with additional data

In this set of experiments we compare models trained on just image data with model trained on image data and additional patient attributes like age, gender and visual acuity. Fig. 5 shows the effect of additional variables on overall accuracy per patient or exact matches, and the F1 score for each pathology.

**Figure 5:**
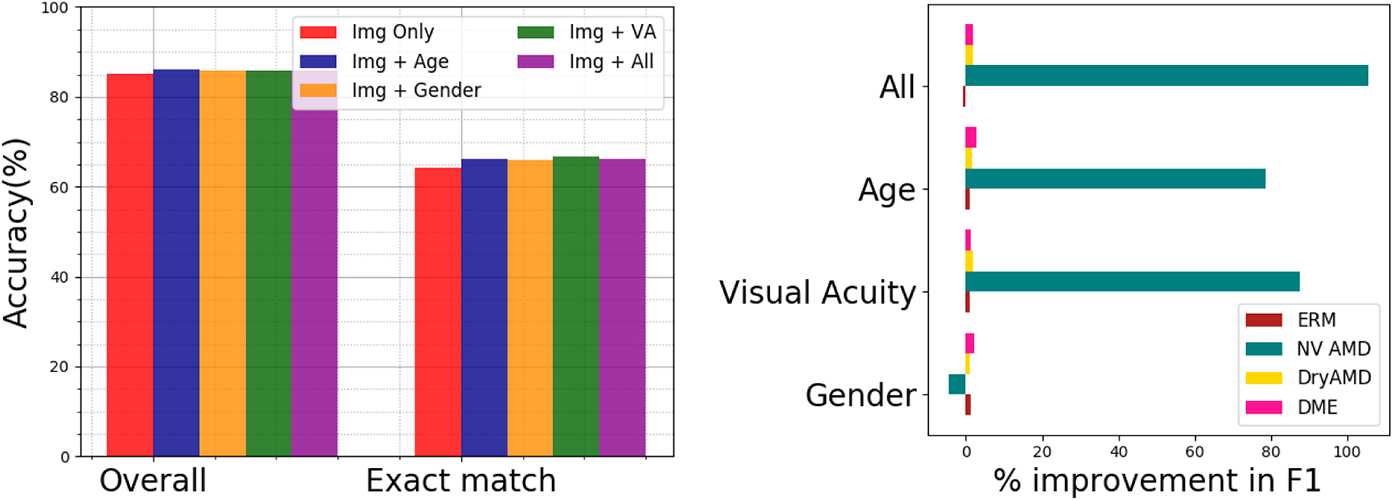
Overall and exact match accuracy (left), improvement in F1 score for per pathology (right) with additional attributes.

### 4.4 Model visualization

We use gradient-weighted Class Activation Mapping (Grad-CAM) and guided Grad-CAM (Selvaraju et al., 2016) to interpret what the models are learning. Grad-CAM uses the class-specific gradient information flowing into the final convolutional layer of a CNN to produce a coarse localization map of the important regions in the image. Guided Grad-CAM fuses Guided Backpropagation (Springenberg et al., 2014) and Grad-CAM visualizations via pointwise multiplication to product visualization that are higher resolution and class discriminative. We show Grad-CAM, guided Grad-CAM for a sample of images from four classes in Fig. 6. The images highlight the regions that the neural network considers important for the diagnosis. T

**Figure 6:**
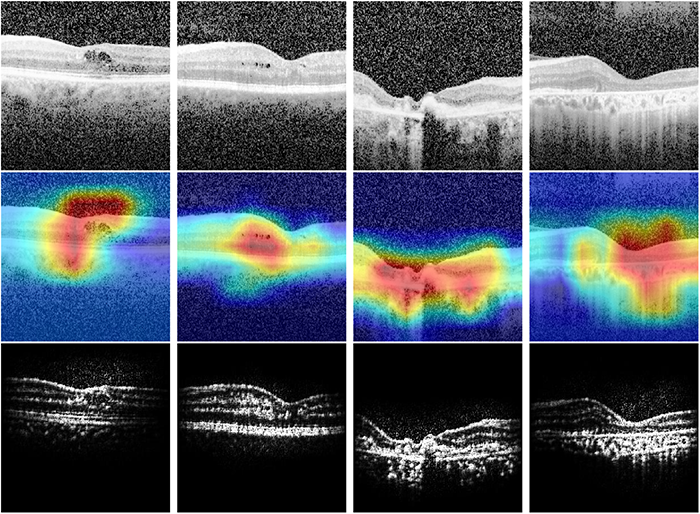
This figure shows what the DNN is learning. The top row is the original OCT scan, central row is the gradient-weighted class activation map and the bottom row is the guided back propagation image overlaid with gradient based class activation map. First column has diagnosis of DME, second ERM and third DryAMD and the fourth NVAMD

## 5. Discussion

In this study the data we used was scraped from EMR, all images with the relevant data were included. No images were excluded because of image quality reasons. Additionally the labels for training were also scraped from the ERM. There were 30 doctors responsible for diagnosing and entering diagnosis related notes, which were used to create true labels. There was no pre-selection done on the images other than selecting middle 11 slices per examination. We did not perform any additional clinical studies to ensure that the diagnosis were consistent. Despite no addition curation of the data we were able to achive overall acurracy of 86% and exact match of 66.7% on the hold out test set of 12,581 images. Since most of the other studies ether manually curated the training data and label Kermany et al. (2018), or performed binary classification Lee et al. (2016)our results are encouraging. Class activated maps and guided class activation maps techiniques were used to determine what the network was learning. The resulting visualizations are consistent with what a clinician would look at when making the diagnosis.

We added other relevant patient information which is often available to clinicians diagosing pathologies on OCT scan such as age, gender and visual acuity. We found that augmenting the deep neural network (DNN) with these attributes was helpful in increasing the F1 score of the pathology which was most often mis-diagnosed by the network. Specifically, F1 score for NVAMD increased by over 100% when providing additional attributes to the DNN. The addition of age and visual acuity was most helpful in boosting the F1 scores while addition of gender information did not result in deterioration of the scores. The impact on the overall accuracy was not as high with this addition as test as well as training samples had the fewest images exhibiting NVAMD. This result shows that augmentation of DNNs with additional relevant attributes of patients when creating computer aided diagnosis sytems can result in higher precision and recall.

We found that transfer learning did not perform well for OCT data in this case. Most transfer learning is done on models pre-trained on three channel images (like ImageNet). With the lower level weights frozen the filters detecting colors resulted in dead features which resulted in loss values to plateau. This resulted in lower accuracy from these models. The training of these model(s) was only done on images from a single academic center and the external generalizability is unknown.

## 6. Conclusion

In this work, we propose using multilabel multiclass classification for OCT retinal images to diganose patients who may exhibit multiple pathologies. To our knowlegde this is the first study on classifying OCT images wih multiple labels. We achieve overall acurracy of 86% and exact match of 66.7%. We show that augmentation of the model with additional patient attributes results in higher F1 scores.

## 7. Ackowledgements

This work is supported in part by NSF grant AITF 1535565 and a gift from Intel, National Eye Institute [K23EY024921]; the Research to Prevent Blindness, New York, NY. Ariel Rokem was funded through a grant from the Gordon and Betty Moore Foundation and the Alfred P. Sloan Foundation to the University of Washington eScience Institute Data Science Environment.

## Footnotes

1 https://research.googleblog.com/2016/08/improving-inception-and-image.html

## References

Pytorch, deep learning framework. URL https://github.com/pytorch.

Yoshua Bengio and Others. Deep learning of representations for unsupervised and transfer learning. ICML Unsupervised and Transfer Learning, 27:17–36, 2012.

Joon Yul Choi. et al. Multi-categorical deep learning neural network to classify retinal images: A pilot study employing small database. PLOS ONE, 12(11), 11 2017.

Jeff Donahue, Yangqing Jia, Oriol Vinyals, Judy Hoffman, Ning Zhang, Eric Tzeng, and Trevor Darrell. Decaf: A deep convolutional activation feature for generic visual recognition. In ICML 2014, pages 647–655, 2014.

Andre Esteva. et al. Dermatologist-level classification of skin cancer with deep neural networks. 542, 2017.

Daniel S. Kermany, Michael Goldbaum, Wenjia Cai, Carolina C.S. Valentim, Huiying Liang, Sally L. Baxter, and Kang Zhang et al. Identifying medical diagnoses and treatable diseases by image-based deep learning. Cell, 172, 2018.

Alex Krizhevsky, Ilya Sutskever, and Geoffrey E. Hinton. Imagenet classification with deep convolutional neural networks. In NIPS 2012, pages 1097–1105. 2012.

Yann LeCun, Yoshua Bengio, and Geoffrey E. Hinton. Deep learning. Nature, 521(7553): 436–444, 2015.

C. S. Lee, D. M. Baughman, and A. Y. Lee. Deep learning is effective for the classification of OCT images of normal versus Age-related Macular Degeneration. ArXiv e-prints, 2016. logMAR. https://en.wikipedia.org/wiki/LogMAR_chart, accessed,2018.

Ariel Rokem. et al. Assessment of the need for separate test set and number of medical images necessary for deep learning: a sub-sampling study. bioRxiv, 2017.

Ramprasaath R. Selvaraju, Abhishek Das, Ramakrishna Vedantam, Michael Cogswell, Devi Parikh, and Dhruv Batra. Grad-cam: Why did you say that? visual explanations from deep networks via gradient-based localization. CoRR, abs/1610.02391, 2016.

Jost Tobias Springenberg, Alexey Dosovitskiy, Thomas Brox, and Martin A. Riedmiller. Striving for simplicity: The all convolutional net. CoRR, abs/1412.6806, 2014.

Eric A. Swanson and James G. Fujimoto. The ecosystem that powered the translation of oct from fundamental research to clinical and commercial impact. Biomed. Opt. Express, 8(3):1638–1664, 2017.

C. Szegedy. et al. Inception-v4, Inception-ResNet and the Impact of Residual Connections on Learning. ArXiv e-prints, 2016.

T. Tieleman and G. Hinton. Lecture 6.5-rmsprop: Divide the gradient by a running average of its recent magnitude, 2012.

Jason Yosinski, Jeff Clune, Yoshua Bengio, and Hod Lipson. How transferable are features in deep neural networks? In NIPS 2014, pages 3320–3328, 2014.

